# Homing in on the rare virosphere reveals the native host of giant viruses

**DOI:** 10.1101/2023.06.27.546645

**Authors:** Amir Fromm, Gur Hevroni, Flora Vincent, Daniella Schatz, Carolina A. Martinez-Gutierrez, Frank O. Aylward, Assaf Vardi

**Affiliations:** Department of Plant and Environmental Sciences, Weizmann Institute of Science, 7610001, Rehovot, Israel; Department of Biological Sciences, Virginia Tech, Blacksburg, Virginia, USA; Google Geo, Israel; Developmental Biology Unit, European Molecular Biological Laboratory, 69117, Heidelberg, Germany; Center for Emerging, Zoonotic, and Arthropod-Borne Pathogens, Virginia Tech, Blacksburg, Virginia, USA 24061

## Abstract

Giant viruses (phylum Nucleocytoviricota) are globally distributed in aquatic ecosystems^1,2^. They play major roles as evolutionary drivers of eukaryotic plankton^3^ and regulators of global biogeochemical cycles^4^. Recent metagenomic studies have significantly expanded the known diversity of marine giant viruses^1,5–7^, but we still lack fundamental knowledge about their native hosts, thereby hindering our understanding of their lifecycle and ecological importance. Here, we aim to discover the native hosts of giant viruses using a novel, sensitive single-cell metatranscriptomic approach. By applying this approach to natural plankton communities, we unraveled an active viral infection of several giant viruses, from multiple lineages, and identified their native hosts. We identify a rare lineage of giant virus (Imitervirales-07) infecting a minute population of protists (class Katablepharidaceae) and revealed the prevalence of highly expressed viral-encoded cell-fate regulation genes in infected cells. Further examination of this host-virus dynamics in a temporal resolution suggested this giant virus controls its host population demise. Our results demonstrate how single-cell metatranscriptomics is a sensitive approach for pairing viruses with their authentic hosts and studying their ecological significance in a culture-independent manner in the marine environment.

## Introduction

Giant viruses commonly referred to as Nucleocytoplasmic large DNA viruses (NCLDV), are abundant and have a broad phylogenetic diversity in aquatic ecosystems^1,2,8^. Infection by giant viruses can have a profound metabolic consequence on the host due to various auxiliary metabolic pathways involved in nutrient uptake, lipid metabolism, and even energy production, that are encoded by the viruses^9^. Some giant viruses infect and lyse bloom-forming alga and thereby play an important role in recycling major nutrients and enhancing the metabolic flux that fuels the ocean microbiome^4^. Moreover, the evolutionary arms race between giant viruses and their hosts can have important consequences for gene transfer^10^ and may even lead to an integration of giant virus genomes into those of their hosts, resulting in dramatic evolutionary consequences^3,11^.

Considering the key ecological role of giant viruses in the ocean, extensive efforts have been made to catalog their diversity across diverse ecosystems worldwide^1,5–7^. Thus, our current knowledge about the ecological diversity of giant viruses stems mainly from metagenomics studies that are conducted on the bulk population level. Furthermore, host-giant virus models in the lab mainly consist of protists (i.e., amoeba) that can phagocytose giant viruses without being their native hosts^12^. Consequently, knowledge about the interactions of giant viruses with their native host is currently limited to only a few model systems, and a deeper understanding of their life cycle and impact on the aquatic environment remains elusive. Current approaches to predict host-virus pairs include computationally correlating between viruses and hosts in metagenomic data^13,14^, finding homology between horizontally transferred genes in the host or viral genomes^1^, and single-cell genomics to detect host and virus DNA in the same cell^15^. Despite these attempts, most metagenome-derived giant viruses still lack reliable predictions of their native hosts. Notably, even single-cell genomics cannot confidently link viruses to their specific hosts because many heterotrophic protists feed on a wide range of viruses in the environment, leading to the detection of viral DNA that is derived from ingestion rather than infection of the native host cells^16–18^.

Single-cell transcriptomics is an innovative approach to tracking host-virus dynamics by detecting the co-expression of a virus and its host transcriptomes within individual cells^19,20^. This sensitive approach enables the detection of active viral infection even in rare viruses that would otherwise be difficult to detect through conventional analysis of bulk metagenomic or metatranscriptomic data ^19,20^.

Here, we develop a single-cell metatranscriptomic approach to map specific viral gene markers to host cells across tens of thousands of single-cell transcriptomes using samples collected in natural planktonic communities. Using this method, we found dozens of infected cells representing eight distinct pairs of hosts and viruses and unraveled the hosts of several giant viruses from multiple lineages, even when the host comprises less than half a percent of the protist community. Overall, single-cell metatranscriptomics provides a sensitive tool for the identification of the native host of giant viruses and tracking their dynamics in the natural environment.

## Results and discussion

To identify novel host-virus interactions in the ocean, we sampled natural plankton communities from an induced *Emiliania huxleyi* bloom during a mesocosm experiment in the Raunefjorden fjord near Bergen, Norway, in May 2018^21^. During this experiment, seven bags were filled with fjord water and monitored for plankton succession for 24 days. Ten samples at a size fraction of 3-20 µm were obtained from four of these bags and fixed on-site, before being processed in the lab (Fig 1a, b). Samples were sequenced using 10X Genomics single-cell (SC) RNA-seq (Fig. 1c), so that cells were barcoded, and each transcript received a unique molecular identifier (UMI). We then mapped the transcripts to a reference database of genes that are conserved in NCLDV and are considered giant virus marker genes, such as viral DNA polymerase beta (PolB), viral type II topoisomerase (TopoII), and major capsid protein (MCP). The expression of these viral marker genes was quantified (Fig. 1d, see Methods for details) and cells with high viral expression were selected (Fig. 1e). Reads from these selected cells were assembled to recover longer transcripts (Fig. 1f, g). To predict the host, we used sequence homology of assembled transcripts to the 18S rRNA gene (Fig. 1h, i). To identify which viruses are infecting these host cells, reads from selected cells were mapped to the database of viral marker genes (Fig. 1h, i).

**Figure 1.**
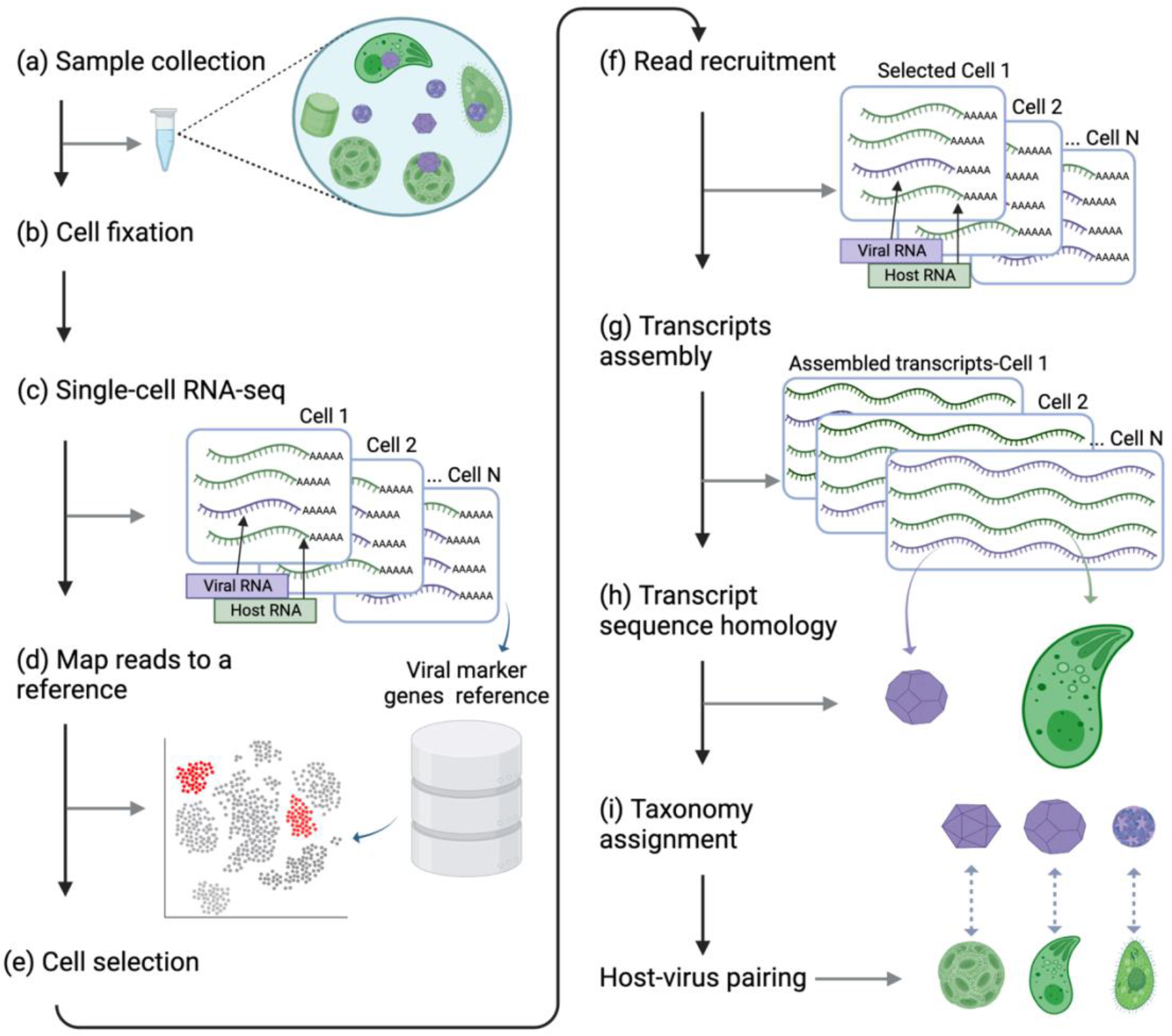
A pipeline for detecting novel host-virus pairs in the natural environment. Black arrows indicate the direction of the pipeline. Gray arrows point to the intermediate output of each step. **(a, b)** Samples were collected from the natural environment and fixed before sequencing. **(c)** Dual SC RNA-seq was performed using the 10X Genomics platform. **(d)** Cells were mapped to a reference of giant virus marker genes to identify cells expressing viral transcripts. **(e)** A subset of cells with high expression of viral transcripts was selected for subsequent analysis. **(f)** Single-cell transcripts were recruited from each selected cell. **(g)** Trimmed single-cell reads (60bp) were assembled to generate longer single-cell transcripts (110-2050 bp). **(h)** The organism encoding the transcript was determined using assembled sequence homology analysis. The virus is identified using raw reads homology. **(i)** Taxonomy was assigned to the host and virus using transcripts and reads from each cell and phylogenetic analysis of 18S rRNA genes (host) and NCLDV marker genes (virus). The figure is created with BioRender.com.

Following this workflow, a total of 972 cells were defined as ‘highly infected’ (see methods; Supplementary Data 1). Most of these cells (n = 754) were infected by *E. huxleyi* virus (EhV), in comparison to 218 cells that were infected by other viruses, confirming the prevalence of infected *E. huxleyi* cells during bloom demise^22,23^. We successfully detect *E. huxleyi*–EhV pairs, confirming this pipeline can be used to detect authentic hosts infected by a well-characterized giant virus. We have previously analyzed in depth the population dynamics of *E. huxleyi* and its virus in this bloom^23^. Here, we wish to identify novel host-virus pairings, thus focusing on cells that were not infected by EhV.

### Novel host-virus interactions at a single-cell resolution

Out of the 218 ‘highly infected’ cells identified, 75 host-virus pairs were defined at the class or division level for the host, and the family level for the virus (Fig. 2, Supplementary Data 2). Viral genes were expressed in protists belonging to diverse and ecologically important taxa such as Chrysophyceae (29%), Prymnesiophyceae (21%), and Dinoflagellata (multiple classes, 9%), as well as the understudied class of Katablepharidaceae (13%) (Fig 2). In about half (56%) of infected cells, viral reads were mapped to multiple families rather than a specific match to one virus lineage. This may imply that the specific virus infecting this host is yet to be discovered and is missing in the reference database. Alternatively, it may also suggest that a single cell can be infected by more than one lineage of viruses, which is known as superinfection^24^. In 44% of the cells (n = 33), at least 90% of viral reads were mapped to a specific virus family (Fig 2). These infected cells represent distinct pairs between eight protist taxa and giant viruses from the order Imitervirales (IM): Mesomimiviridae (IM_01), Mimiviridae (IM_16), IM_09, and IM_07. This is consistent with the reported dominance of the *Imitervirales* in marine ecosystems^2^.

**Figure 2.**
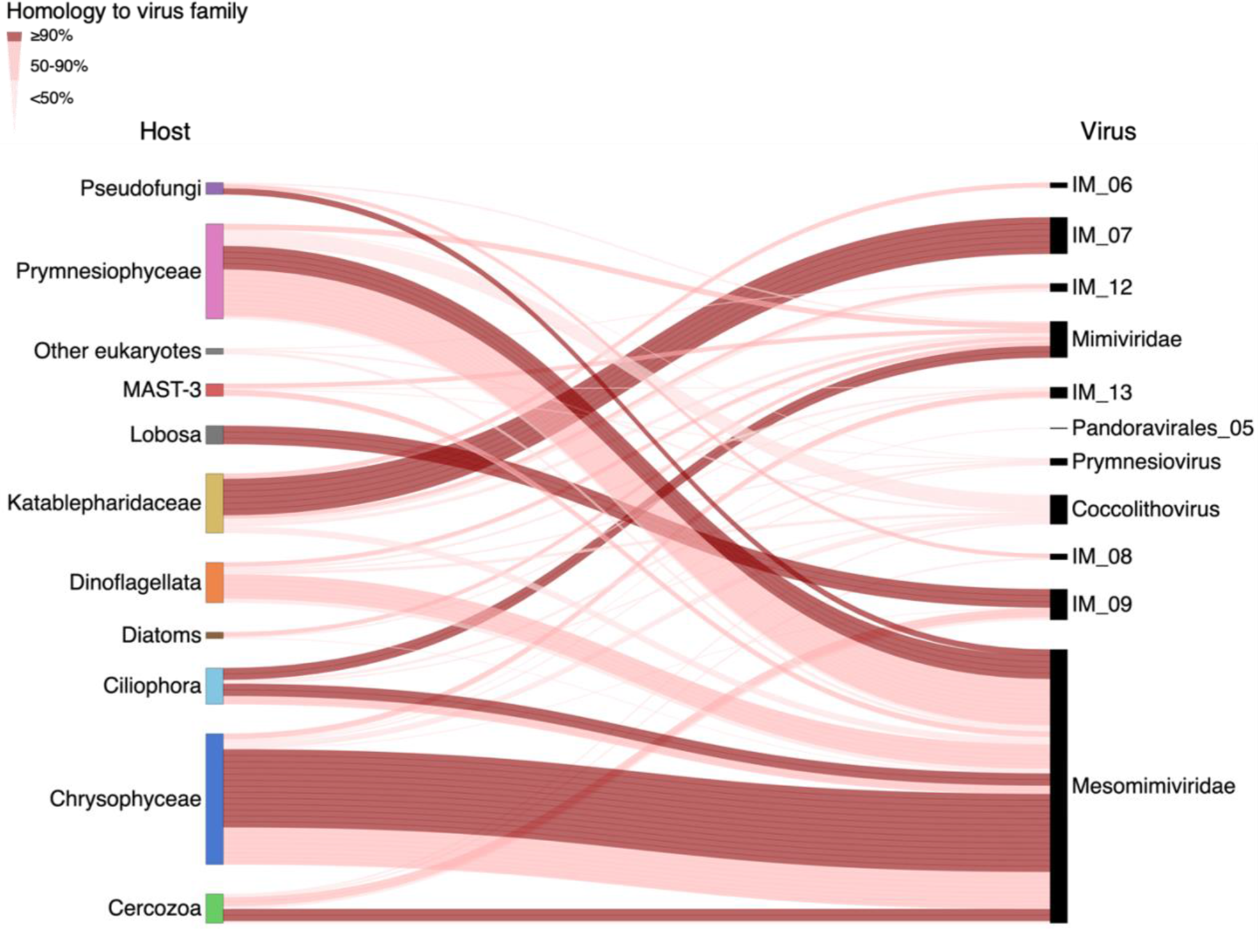
Assigning host cells and their actively infecting giant viruses within the algal bloom. Connecting lines represent the predicted pairing of host cells to their infecting viruses (n=75). Dark, thick lines represent unambiguous pairing (≥90% of viral reads, 33 cells) of a virus family to its specific designated host. Thinner, lighter lines represent less specific pairing (<90% of viral reads, 39 cells).

Out of the identified virus families, Mesomimiviridae was the most prevalent group of viruses actively infecting the cells (66% of distinct links, 22 cells), and they infect multiple cells from various families, mostly Chrysophyceae (13 cells) and Prymnesiophyceae (4 cells). This is consistent with the observation that giant viruses from the Mesomimiviridae are the most abundant and widespread family of giant viruses in the ocean^8^, and are known to infect bloom-forming Prymnesiophyceae algae as *Chrysochromulina* and *Phaeocystis*^25,26^.

In addition to the known association between Mesomimiviridae viruses and Prymnesiophyceae hosts, we identify seven completely novel host-virus pairs which represent the first description of their interaction *in situ*. For instance, the IM_9 viral family was found to actively infect cells of Lobosa (Amoebozoa, 3 cells), and Mimiviridae (2 cells). As for the virus family IM_07, all 6 infected cells were identified as Katablepharidaceae, heterotrophic flagellates related to Cryptophytes^27^. So far, no nuclear genome of any Katablepharidaceae has been sequenced, and no virus was described to infect this class. To our knowledge, this is the first report of a specific host infected by an IM_07 virus. Only 19 metagenome-assembled genomes (MAGs) from this viral group are currently available, and all have been found in aquatic ecosystems^8^. To verify the phylogenetic position of the newly identified host-virus pairs, we constructed phylogenetic trees from transcripts assembled from both host cells and their infecting viruses (Extended Data Fig. 1). We thus elucidated multiple connections, including previously unknown interactions, between important protist classes and a variety of virus families. Future research should explore if newly identified viruses, that are currently absent from our database, can be detected in these samples.

### Tracking multiple viral infections co-occurring in a natural population

To examine co-occurring viral infections in different protist populations during bloom succession, all reads were mapped to a custom host-virus reference database which represents the different protist groups in the population. This database was generated based on the single-cell transcriptomes of the selected infected cells (Fig. 2). To this host-virus reference, we added genes from EhV and *E. huxleyi*, which dominated the bloom^22^. Data derived from a total of 18,793 RNA-sequenced single cells were mapped to the host-virus reference and visualized using Uniform Manifold Approximation and Projection (UMAP) representation. Each cell was assigned taxonomy by 18S rRNA homology (Fig. 3a). As expected, the largest cluster was dominated by Prymnesiophyceae, the class of *E. huxleyi*, the dominant species in the bloom^21,23^. A smaller, yet distinct cluster consists of mainly MArine STramenopiles (MAST)-3, a group suggested to be one of the most abundant MAST groups in the ocean^28^ (Fig. 3a, in red). A secondary cluster consists of diverse assemblages and includes 81 infected cells, identified by highly expressing contigs of viral origin (Fig. 3b, see methods). These cells belong to previously described taxa (Fig 2): Pseudofungi, Dinoflagellata, Chrysophyceae, Prymnesiophyceae, and Katablepharidaceae (Fig. 3b). This analysis revealed active viral infection at a single-cell level, occurring in different protist host cells originating from diverse taxa in the natural environment. This hidden host-virus dynamics and diversity are often completely masked by a viral infection of the dominated bloom-forming algae. Therefore, single-cell metatranscriptomics provides a new opportunity to detect active viral infection at the cellular level and provides a sensitive lens into host-virus dynamics in the rare virosphere and its possible ecological implications.

**Figure 3.**
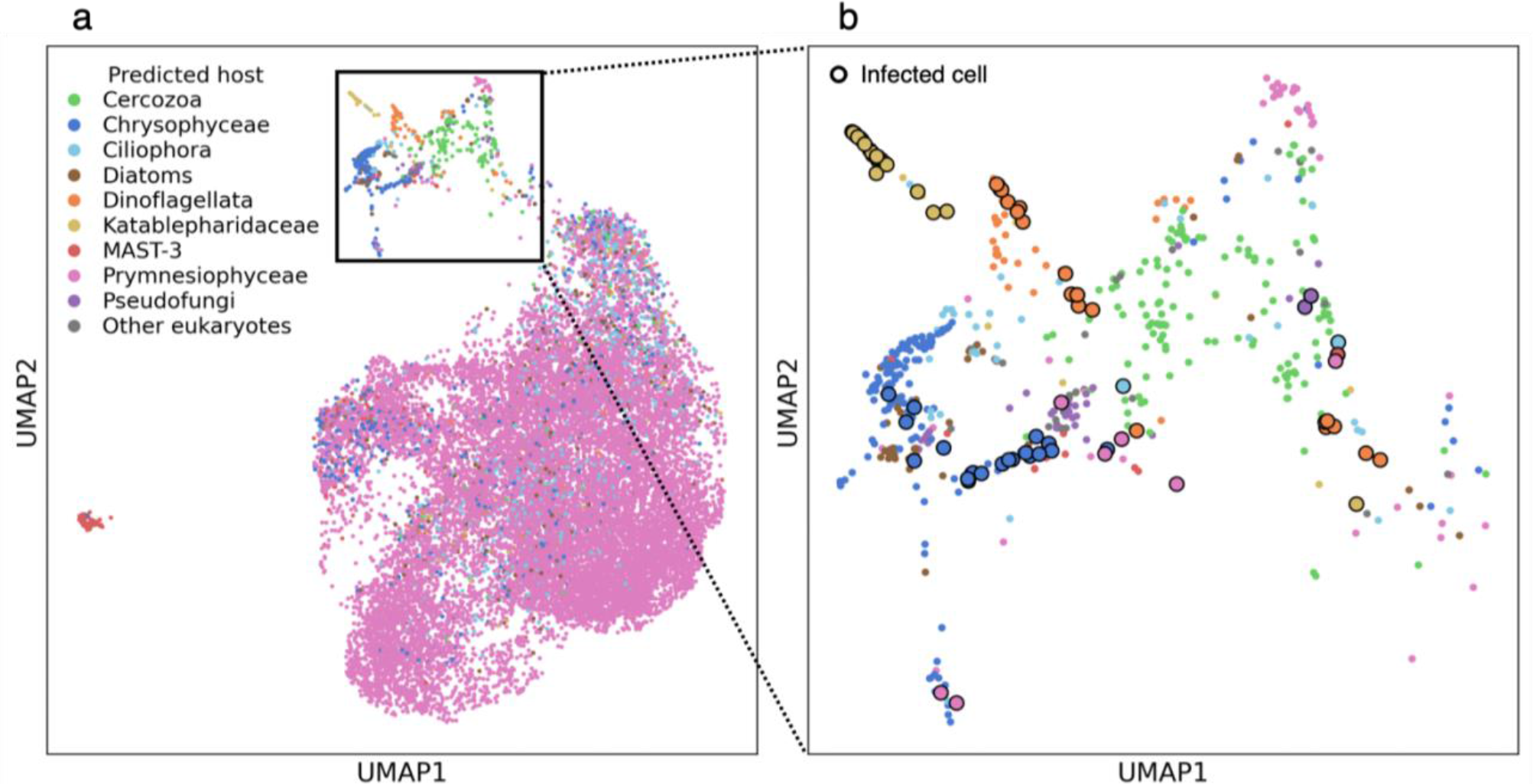
Co-occurring viral infection of diverse protist groups at a single cell resolution. A UMAP projection of cells that were mapped to the host-virus reference of single-cell transcripts and EhV-*E. huxleyi* genes (n=18,793). Each cell is colored according to its predicted taxonomy. The black square marks the subset of cells that contains the highest diversity of taxa **b.** A close-up view on viral infection in a subset of cells from diverse taxa (n=612). Cells that express at least 10 UMIs of viral transcripts are enlarged and marked with a black edge (n=81).

### Identifying host-virus interactions in rare protists from the Katablepharidaceae class

Our approach enables mapping active infection events among diverse protist host cells and can provide a sensitive means to detect rare infected cells. As a case study, we tracked Katablepharidaceae cells for which we detected infection by giant viruses of the IM_07 family (Fig 2, Fig. 3b). Katablepharidaceae represent less than 0.5% of all detected cells (Fig 3a, n=70 cells out of 18,793 cells), and about 6% of the population in the diverse cluster (Fig 3b, n=37 cells out of 612 cells). In this cluster, a distinctive subpopulation of highly infected Katablepharidaceae cells could be observed that makes up about a third of the infected cells in this cluster (Fig. 3b, n=27 cells out of 81 infected cells). To further explore this infected subpopulation, we pulled together and assembled the transcriptome from 26 infected Katablepharidaceae cells from the same sample (bag #4, day 20 of the mesocosm experiment). These assembled contigs matched the 18S rRNA gene of *Leucocryptos marina* (>95% identity, E-value ≤ 10^-10^, Supplementary Data 3). The best match for the virus infecting Katablepharidaceae cells is the IM_07 member GVMAG-M-3300020187-27 (identity ≥ 99%, E-value ≤ 10^-10^), a virus that was first assembled from a metagenomic analysis on samples obtained from Kabeltonne, Helgoland, North Sea, but has never been isolated^1^. This is the first virus that is identified to infect the genus *Leucocryptos* and the class Katablepharidaceae in general, and the first definitive host for giant viruses of the IM07 lineage.

### Genomic assembly and characterization of the predicted *Leucocryptos* virus

Katablepharidaceae are a class of flagellated heterotrophic plankton, that consists of 5 species, none of which have a published nuclear genome^29^. *Leucocryptos marina*, the closest relative to the predicted host, is a phagotrophic heterotroph abundant in coastal waters with high plankton productivity^30^. The predicted *Leucocryptos* virus has the largest genome that has been recovered from the IM_07 lineage (950 kbp), and it encodes for 894 genes (Fig. 4a)^1,8^. Reads from the infected cells were mapped to the assembled viral genome and the expression of viral genes was examined (Fig 4a, b, Supplementary Data 4). Viral transcription varies greatly between predicted contigs, with some contigs showing no expression at all, which might be the result of a binning error during the original metagenomic assembly. The virus encodes a complex repertoire of 13 proteins likely involved in manipulating cellular stress responses and cell fate regulation, including a predicted Bax-1 apoptosis inhibitor, a metacaspase homolog, a homolog of heat-shock protein 90 (HSP90), two homologs each of HSP70, and 8 homologs of DnaJ (HSP40) genes. These proteins are placed inside well-defined viral clades separated from eukaryotic clades and together with other viruses of the *Imitervirales* group (Extended Data Fig. 2), suggesting that these genes were horizontally transferred from host genomes to viral genomes early in their evolution ^3^. HSP90 and HSP70 are also among the most highly expressed viral genes (Fig 4b, Supplementary Data 4). Heat shock proteins play a role in the life cycle of many viruses, mostly in viral replication, and in some cases are encoded by the virus^31^. Heat shock proteins of the Herpes Simplex virus were shown to regulate virus-induced apoptosis and regulate other heat shock proteins^32^. In addition, viral-encoded metacaspases have been suggested to regulate host cell death and were identified in diverse giant viruses from the marine environment^33,34^. The high prevalence and expression of these cell fate regulators encoded by the *Leucocryptos* virus, suggests that they have an important function in its life cycle by control of their host’s cell death. Interestingly, the virus also encodes for 9 predicted major capsid (MCP) proteins, a high number even for a giant virus (Fig. 4b)^1^. Some of these predicted MCPs are co-localized in the genome, suggesting gene duplication. Unlike the anti-apoptotic genes, the MCP genes are lowly expressed in SC-RNAseq analysis. This pattern of expression may suggest that infection is at an early phase.

**Figure 4.**
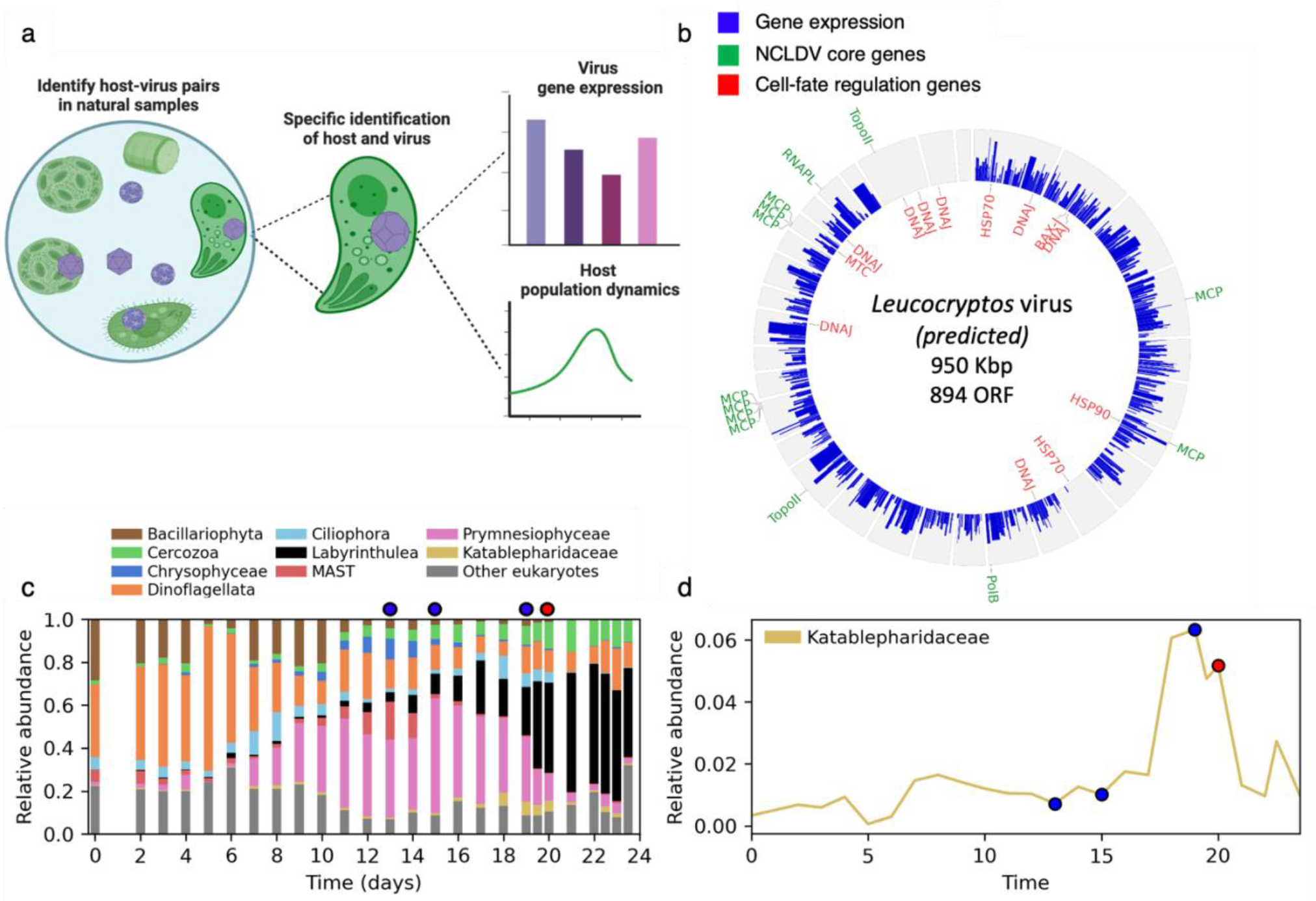
Characterization of *Leucocryptos* virus genome and its suggested impact on the population dynamic of Katablepharidaceae. **a.** A graphic scheme of the *Leucocryptos* virus characterization, from its detection in single cells to finding its putative host and eventually tracking its gene expression and host population dynamics in natural populations. **b.** A Circos plot of the predicted *Leucocryptos* virus. Gene expression (expected read counts, log2 transformed) from 26 infected cells identified as *Leucocryptos* is shown in the blue bars. Key predicted genes are shown in their corresponding location. In green, core NCLDV genes: PolB: DNA polymerase beta; TopoII: Type II topoisomerase; RNAPL: RNA polymerase; MCP: Major capsid protein; PolB: DNA Polymerase subunit. In red, cell-fate regulation genes: HSP: Heat-shock protein; DNAJ: DnaJ chaperone; BAX-I: Apoptosis regulator BAX inhibitor; MTC: Metacaspase. The genome of the virus is shown as circular for convenience. **c.** Relative abundance of different taxonomic groups in bag 4 during the mesocosm experiment. Samples for SC-RNAseq were collected on marked days: 13,15,19 and 20. The red marker points to the time when most infected cells were detected. **d.** Relative abundance of the Katablepharidaceae class in bag 4 during the time course of the mesocosm experiment. Markers are as in panel 4b.

### Population dynamics of Katablepharidaceae following a viral infection

Upon pairing a specific giant virus to its host, we track host-virus dynamics during algal bloom succession in the mesocosm experiment (Fig 4a). The relative abundance of Katablepharidaceae was monitored using amplicon sequence variants (ASVs) of 18S rDNA^21^ and compared to different taxonomic classes that were detected in the SC-RNAseq (Fig. 4c). These results reveal the heterogeneity of the population and confirm the presence of the classes that were found on the single-cell level using SC-metatranscriptomics. The ASV analysis confirmed the rarity of Katablepharidaceae in the community compared to other taxa analyzed here, such as Prymnesiophyceae, Dinoflagellata, and Cercozoa. Interestingly, the Katablepharidaceae class increased in abundance from 1% of the community on day 15 to a maximum of 6% on day 19, followed by a population decline back to 1% only two days later, on day 21 (Fig. 4d). This sharp demise in the relative abundance of Katablepharidaceae was observed after day 20, the same day in which we detected that 86% (n=26) of the observed Katablepharidaceae single cells were infected. This strongly suggests that the IM07 lineage virus was responsible for the population’s demise. The decrease in the population coincides with a sharp increase in the population of Labyrinthulea, which was suggested to act as decomposers during bloom demise^21^. If these protists benefited from decomposing the infected *E. huxleyi* cells, it is also possible that they similarly benefitted from decomposing the infected *Leucocryptos*.

### Conclusions

Research in the last decade has revealed that giant viruses are ubiquitous components of ecosystems around the globe^1,2^. The diversity of numerous giant viruses particularly in the marine environment has increased in recent years, along with a growing interest in their unique life cycle and ecological role. Still, a major knowledge gap is to identify their still unknown native host cells in order to gain insights into their infection dynamics and ecological impact. Here we show that single-cell metatranscriptomics is a novel highly sensitive approach that can be used to identify the native hosts and the transcriptional activity of these viruses in the natural environment, even if they comprise only a small fraction of the total community. We analyzed samples from algal bloom and linked multiple host cells with their infecting virus at a single-cell resolution, including a rare infection of a giant virus. This finding led to the discovery of the first virus to infect the Katablepharidaceae class and the first suggested host for the recently defined group of giant viruses, the IM_07 virus family^8^. This virus was described before in a metagenomic analysis^1^, and these results demonstrate how our new approach can be utilized to find the hosts of viruses that were described in bulk metagenomic data, and also to explain a population dynamic in an algal bloom^21^. This approach demonstrates how single-cell metatranscriptomics has the potential to connect the wealth of metagenomic data acquired on marine viruses with a mechanistic knowledge of their lifestyle resolved by single-cell transcriptomics.

Better mapping of the rare virosphere can help understand the dynamics of complex ecosystems, as rare species are often more active than abundant species, have a high per-organism contribution to the community metabolic potential, and enhance the functionality of abundant species^35^. Active infection in rare virosphere can serve as a seed bank population for the subsequent infection dynamics^36^, especially during the phase of post-bloom demise as in our study, in which the dominance of available host cells is rapidly shifting in genetic composition. Moreover, recent attempts to quantify viral infection rates have shown how low levels of infection are common in marine ecosystems and may have important consequences for viral persistence over broad geographic areas^37,38^. Tracking active viral infection using single-cell approaches may provide insights into the ecological importance of viruses in the marine environment and will help to bridge the gap between large-scale environmental metatranscriptomics and the mechanistic view of the life cycle of diverse protist host cells.

## Methods

### Mesocosm core setup and sampling procedure

Samples were obtained during the AQUACOSM VIMS-Ehux mesocosm experiment in Raunefjorden near Bergen, Norway (60°16′11N; 5°13′07E), in May 2018. Seven bags were filled with 11m^3^ water from the fjord, containing natural plankton communities. Algal blooms were induced by nutrient addition and monitored for 24 days as previously described^21^. 10 samples were collected from four bags, as follows: From bag 3, on days 15 and 20 (named B3T15, B3T20 correspondingly). From bag 4, on days 13, 15,19, and 20 (named B4T13, B4T15, B4T19, and B4T20, correspondingly). From bag 6, on day 17 (named B6T17). From bag 7, on days 16, 17, and 18 (named B7T16, B7T17, and B7T18, correspondingly).

Samples were initially filtered as follows: 2 liters of water were filtered with a 20 µm mesh and collected in a glass bottle. The cells were then concentrated through gentle gravity filtration on a 3 µm polycarbonate filter (Whatman), mounted on a reusable bottle top filter holder (Thermo Fischer). The biomass on the filter was regularly resuspended by gentle pipetting.

For samples B7T16, B7T18, B4T15, B3T15, B6T17, B7T17, and B4T19 the 2 liters of seawater were concentrated down to 100 ml, distributed in two 50 ml tubes, which corresponds to a 200 times concentration. For B4T13, the concentration factor was 140 times. For B4T20 and B3T20, the concentration factor was 100 times. The different concentration factors are explained by filter clogging and various field constraints including time of processing. For all samples except B3T20, the 50 ml tubes were centrifuged for 4 min at 2500g, after which the supernatant was discarded. Pellets corresponding to the same day and same bag were pooled and resuspended in a final volume of 200 µl of chilled PBS. 1800 µl of pre-chilled high-performance liquid chromatography (HPLC) grade 100% methanol was added drop by drop to the concentrated biomass. For B3T20, the concentrated biomass was centrifuged for 4 min at 2500g, resuspended in 100 µl of chilled PBS, to which 900 µl of chilled HPLC grade 100% methanol was added. Then, samples were incubated for 15 min on ice, and stored at -80°C until further analysis.

### Library preparation and RNA-seq sequencing using 10X Genomics

For analysis by 10X Genomics, tubes were defrosted and gently mixed, and 1.7 ml of the samples were transferred into an Eppendorf Lowbind tube and centrifuged at 4°C for 3 min at 3000g. The PBS/methanol mix was discarded and replaced by 400 µl of PBS. Cell concentration was measured using an iCyt Eclipse flow cytometer (SONY), based on forward scatter. Cell concentration ranged from 1044 cells ml^-1^ to 9855 cells ml^-1^. All concentrations were brought to 1000 cells ml^-1^ to target 7000 cells recovery, according to the 10X Genomics Cell Suspension Volume Calculator Table provided in the user guide. Two libraries were prepared and sequenced on different occasions: B4T19 and B7T17 in January 2020 and B3T15, B3T20, B4T13, B4T15, B4T20, B6T17, B7T16, and B7T18 in August 2020. All the following steps for library preparation were accomplished according to the manufacturer’s protocol for Chromium Next GEM Single Cell 3’ Reagent Kit v3.1. Libraries were sequenced using NextSeq^®^ 500 High Output kit (75 cycles).

Raw 10X reads were trimmed using TrimGalore (v. 0.6.5), a Cutadapt wrapper^39^, and the poly-A tail was removed.

### Detection of infected cells in the single-cell RNA-seq data using a custom viral genes database

To detect viral transcripts, a reference was built from a database of conserved genes in NCLDVs^6^. To remove redundancy, the proteins were clustered using CD-HIT at 90% identity^40^. From this database, a reference was created using the 10X Genomics Cell Ranger mkref command. The Cell Ranger Software Suite (v. 5.0.0) was used to perform barcode processing and single-cell unique molecular identifier (UMI) counting on the fastq files using the count script (default parameters). For downstream analysis, 972 cells were selected as ‘highly infected’ based on the following criteria: (a) expression of more than one viral gene (>1), (b) expression of at least one gene with a UMI count greater than one (>1), and (c) cell expresses in total ≥10 viral UMIs.

### Identifying the taxonomy of individual cells by sequence homology to ribosomal RNA

Raw reads from each cell were pulled by the cell’s unique barcode identifier using seqtk v. 1.2. Reads were then trimmed, and poly-A was removed, using TrimGalore (v. 0.6.5), a Cutadapt wrapper^39^. Trimmed reads from each cell were assembled using rnaSPAdes 3.15^41]^ with kmer sizes: 21,33. Raw reads pulling, trimming, and assembly was wrapped using an in-house script. To identify the taxonomy of the cells, assembled contigs from each cell were matched against 18S rRNA sequences from the Protist Ribosomal Reference (PR2)^42^ and metaPR2^43^. To remove redundancy, the sequences in each of these databases were clustered using CD-HIT v. 4.6.6 at 99% identity^40^. Contigs were filtered using SortMeRNA v. 4.3.6^44^ with default parameters against the PR2 database and then aligned to the PR2 and metaPR2 databases using Blastn^45^, at 99% identity, E-value ≤ 10^-10^ and alignment length of at least 100 bp. Contigs were ranked by their bitscore, and only the best hit was kept for each contig. Each contig was assigned to one of the following taxonomic groups that were prevalent in the sample: the classes Bacillariophyta, Prymnesiophyceae, Chrysophyceae, MAST-3, and Katablepharidaceae, the divisions Pseudofungi, Lobosa (Amoebozoa), Ciliphora (Ciliates), Dinoflagellata and Cercozoa. Contigs that matched other groups were assigned as “other eukaryotes”. Contigs that matched more than one of these taxonomic groups were considered non-specific and were therefore ignored. Cells that transcribe 18S rRNA transcripts homologous to more than one taxonomic group were conservatively omitted. None of the cells that were assigned “other eukaryotes” had contigs with conflicting annotations (contigs matching different classes).

### Identifying the infected virus using a homology search against a custom protein database

To identify transcripts derived from giant viruses, reads from the detected 972 infected cells were compared to a custom protein database using a translated mapping approach. To ensure that as many giant viruses as possible were represented, a database was constructed by combining RefSeq v. 207^46^ with all predicted proteins in the Giant Virus Database^8^. The proteins were then masked with tantan^47^ (using the -p option) and generated the database with the lastdb command (using parameters -c, -p). To identify the infecting virus, the raw sequencing reads in each of the 972 single cells transcriptomes were compared to the constructed database using LASTAL v. 959^48^ (parameters -m 100, -F 15, -u 2) with best matches retained. The same procedure was done for the assembled transcripts from each cell to identify viral transcripts. The results were analyzed at different taxonomic levels, consistent with either the Giant Virus Database (for giant viruses) or NCBI taxonomy^29^ (everything else).

754 Cells whose best matching virus was coccolithovirus were omitted from the downstream analysis since EhV-infected cells were already reported to be abundant in the algal bloom^23^, and our analysis aims to explore other host-virus pairs.

### Plotting host-virus pairs in a Sankey plot for host cells and their infecting giant viruses

Of the 218 cells detected as infected by viruses other than EhV, 75 cells were selected that could be identified using 18S rRNA and have at least 10 reads mapped to one of the virus families (Supplementary Data 2). Only links representing at least 10% of the mapped reads in each cell are shown, in order to highlight the strong links. The Sankey plot was constructed using Holoviews v. 1.15.4.

### Phylogenetic trees of viral and host marker genes

For phylogenetic analysis, 33 cells were chosen based on a strong correlation (≥90% of viral reads mapped to one virus family) between the host and a virus.

To obtain reference 18S rRNA sequences to include in a phylogeny, all transcripts assembled from these cells were compared to the PR2 database^42^ using BLASTN v. 2.9.0+ (parameters - perc_identity 95, -evalue 10^-10^, -max_target_seqs 20, -max_hsps 1). Sequences shorter than 1000 bp were removed from the reference and the remainder of sequences were de-replicated with cd-hit v. 4.7^40^ (-c 0.99) to prevent the inclusion of excessive nearly-identical references. Sequences were aligned with ClustalOmega v. 1.2.3^49^(default parameters), and diagnostic trees were created with FastTree 2.1.10^50^ for quick visualization of trees and for pruning long branches. The final phylogenetic trees were constructed with IQ-TREE v. 2.1.2^51^ (parameters m TEST -bb 1000 -T 6 --runs 10) using ultrafast bootstraps^52^. To identify major capsid protein sequences in the single-cell transcriptomes, proteins were first predicted using FragGeneScanRs v. 1.1.0^53^ (parameters -t, illumina_10). The resulting protein sequences were compared to MCP proteins in the Giant Virus Database with BLASTP v. 2.12.0+ (parameters -evalue 10^-3^, -max_target_seqs 20, -max_hsps 1) as well as to a custom MCP HMM that were previously designed^6^ using hmmsearch in the HMMER3 v. 3.3.2 package^54^ (E-value ≤ 10^-3^). The results of these searches were manually inspected, and sequences were subsequently aligned with ClustalOmega v. 1.2.3 (default parameters). Similarly, as with the 18S rRNA sequences, diagnostic trees were first made with FastTree 2.1.10 and pruned long branches before making a final tree with IQ-TREE v. 2.1.2 (parameters m LG+I+F+G4 -bb 1000 -T 6 --runs 10) using ultrafast bootstraps.

### Single-cell RNA-seq data alignment to a custom reference

A new host-virus reference database was curated from the transcriptome of the infected cells (Fig. 2). Repetitive sequences were removed using BBduk (BBtools 38.90)^55^. An Additional long repetitive sequence was removed manually. A database of *E. huxleyi* and EhV genes, which were shown to be abundant in the samples^23^, was also added to this reference to specifically detect *E. huxleyi* cells and to avoid a non-specific mapping of reads from these cells to other contigs. For EhV, the predicted CDSs in the EhVM1 were used as a reference^56^. For the host, an integrated transcriptome reference of *E. huxleyi* was used as a reference^57^. Viral transcripts in the database were identified using a homology search against a custom protein database as described above. A reference was created from the database using the Cell Ranger mkref command. Fastq files were mapped to this reference database using 10X Genomics Cell Ranger v. 5.0.0 count analysis.

### Preprocessing of transcript abundance and dimensionality reduction

Cells with zero UMIs as well as cells with the lowest 1% number of UMIs were removed for downstream analyses. Cells with the highest 1% number were also removed to prevent cases of doublet or multiplet cells. The raw UMIs of 30,521 cells were further preprocessed using the Python package scprep v. 1.0.10: Low expressing genes were filtered with filter.filter_rare_genes and min_cells=2, expression was normalized by cell library size with normalize.library_size_normalize, and the data was scaled with transform.sqrt.

To represent the cells in two dimensions based on their gene expression profiles, dimensionality reduction using was performed using scprep v. 1.1.0 package PCA (method=’svd’, eps=0.1) and Uniform Manifold Approximation and Projection (UMAP) dimensionality reduction was conducted using the UMAP method in the manifold package of the Python library scikit-learn v. 0.24.1 (minimum distance=0.4 spread=2, number of neighbors=7).

### Assigning taxonomy to each detected cell using rRNA homology search

To identify the taxonomy of each of the detected cells, reads from each cell were assembled independently. The taxonomy of the cells was determined by 18S rRNA homology to one of the following groups, which were abundant in the population: the classes Bacillariophyta, Prymnesiophyceae, Chrysophyceae, MAST-3 and Katablepharidaceae, the divisions Pseudofungi, Ciliphora (Ciliates), Dinoflagellata and Cercozoa. Other taxonomic groups were clustered under “Other eukaryotes”. 18,793 cells were identified this way, and 11,728 cells that could not be identified were excluded from the plot for convenience. Cells with 18S rRNA contigs homologous to more than one taxonomic group were also conservatively omitted. Cells expressing at least 10 viral UMIs as described above were considered infected.

### Identifying the *Leucocryptos* host and its virus using homology search

To better identify the detected Katablepharidaceae cells and to identify their infecting virus, 26 infected Katablepharidaceae cells from bag #4, day 20 were selected. Reads from these cells were retrieved using the unique molecular identifier, and then trimmed using TrimGalore v. 0.6.5, a Cutadapt wrapper^39^. Trimmed reads from all these cells were assembled altogether using rnaSPAdes v. 3.15^41^. To identify the specific Katablepharidaceae host, assembled contigs were matched against the PR2 rRNA database and the using blastn at 90% identity, E-value ≤ 10^-10^, and alignment length ≥ 100bp. Contigs best matched to an unknown Katablepharidaceae (>99% nucleotide identity), but after removing unidentified genera, these contigs best matched (>95% nucleotide identity) the Katablepharidaceae species *Leucocryptos marina*. Transcripts that matched classes other than Katablepharidaceae were matched against the entire NCBI database using the NCBI web server^58^. They too mostly matched Katablepharidaceae genes, specifically 28S rRNA or internal transcribed spacer (ITS) sequences (Supplementary Data 3). To identify the specific infecting virus, transcripts were matched against an NCLDV gene marker database^6^ at 90% identity, E-value ≤ 10^-10^, and alignment length ≥ 100bp. After finding homology to *Leucocryptos* and the virus GVMAG-M-3300020187-27^1^, gene expression was calculated using RSEM v.1.3.1^59^ (parameter --fragment-length-mean 58). The genomic features of the virus were taken from Schulz (2020)^1^ and the viral genome was plotted using ShinyCircos v. 2.0^60^. Gene expression in the plot is measured in expected counts, after log 2 transformation. The relative abundance data in Fig. 4 was obtained from an 18S rRNA amplicon sequencing on a size fraction of 2µm in bag #4 during the mesocosm experiment^21^. In Fig. 4c, relative abundance is calculated per taxa as a fraction of all amplicon sequencing variants (ASV) excluding metazoans. Fig. 4d shows the fraction of Katablepharidaceae out of all ASVs matching Katablepharidaceae (excluding metazoans).

### Phylogenetic tree of Katablepharidaceae ASVs and 18S rRNA genes

To verify the taxonomy of the ASVs, A phylogenetic tree was constructed of 89 ASVs identified as Katablepharidaceae, selected 18S rRNA sequences of Katablepharidaceae and other species from the PR2 database, and the longest single cell assembled contig from the infected Katablepharidaceae cells. Sequences were aligned with ClustalOmega v. 1.2.4 (default parameters). A diagnostic tree was first made with FastTree 2.1.10^50^ for pruning long branches before making the final tree with IQ-TREE^51^. All but three ASVs and one PR2 sequence clustered together with the assembled *Leucocryptos* transcript, verifying the taxonomy of 97% of the ASVs used in the relative abundance analysis (Extended Data Fig. 3).

### Phylogenetic trees of viral heat-shock proteins and metacaspase

To examine the evolutionary history of the heat-shock proteins encoded in GVMAG-M-3300020187-27, phylogenetic trees of these proteins were constructed together with homologs present in eukaryotes, bacteria, archaea, and other giant viruses. For this, a custom database of proteins from reference genomes was compiled from EggNOG v. 5.0^61^ (eukaryotes), bacteria and archaea (the Genome Taxonomy Database v. 95)^62^, and other giant viruses (the Giant Virus Database^8^). For bacterial and archaeal genomes in the GTDB, proteins were predicted first with Prodigal v. 2.6.3^63^ using default parameters. Proteins were searched against Pfam models for each protein using hmmsearch with the noise cutoff (--cut_nc), and subsequently aligned sequences with ClustalOmega v. 1.2.3 (default parameters). Phylogenetic trees were constructed trees using IQ-TREE v. 2.1.2^51^ (parameters m TEST -bb 1000 -T 6 --runs 10) using ultrafast bootstraps and with the best model determined with ModelFinder^64^.

## Supporting information

The homology search results of single-cell reads mapped against the viral gene markers database.

A summary table of host-virus pairs (as seen in Fig. 2)

Blast results of infected Katablepharidaceae subpopulation against the PR2 database and against the viral genes markers database.

Genomic features, gene annotations, and gene expression of the virus GVMAG-M-3300020187-27.

## Acknowledgments

We thank Adva Shemi for her comments and suggestions for this manuscript. This study was supported by the Simons Foundation grant (no. 735079) “Untangling the infection outcome of host-virus dynamics in algal blooms in the ocean” awarded to A.V. and NIH grant (no. 1R35GM147290-01) awarded to F.O.A.

## Author contribution

A.F., G.H., F.O.A. and A.V. designed and conceptualized the project and wrote the paper. A.F. and G.H. designed and wrote the scripts for data analysis. F.V. and D.S. collected the natural samples and prepared the single-cell transcriptomics libraries. F.O.A. and C.A.M.G. conducted phylogenetic analysis and viral homology search. All other data analysis was conducted by A.F. All authors read and edited the manuscript.

These authors contributed equally: A.F and G.H.

## Correspondence

Assaf Vardi

Frank O. Aylward

## Data Availability

Additional data used in this paper, including UMI tables generated from 10X Cell Ranger, extended Blast result tables, assembled transcripts, and other files that can be used to reproduce our results are available at Dryad: https://doi.org/10.5061/dryad.s7h44j1c9.

## Code Availability

All data management and analysis codes are open for review and reuse and archived online at GitHub: https://github.com/vardilab/host-virus-pairing.

## Extended data

**Extended Data Figure 1.**
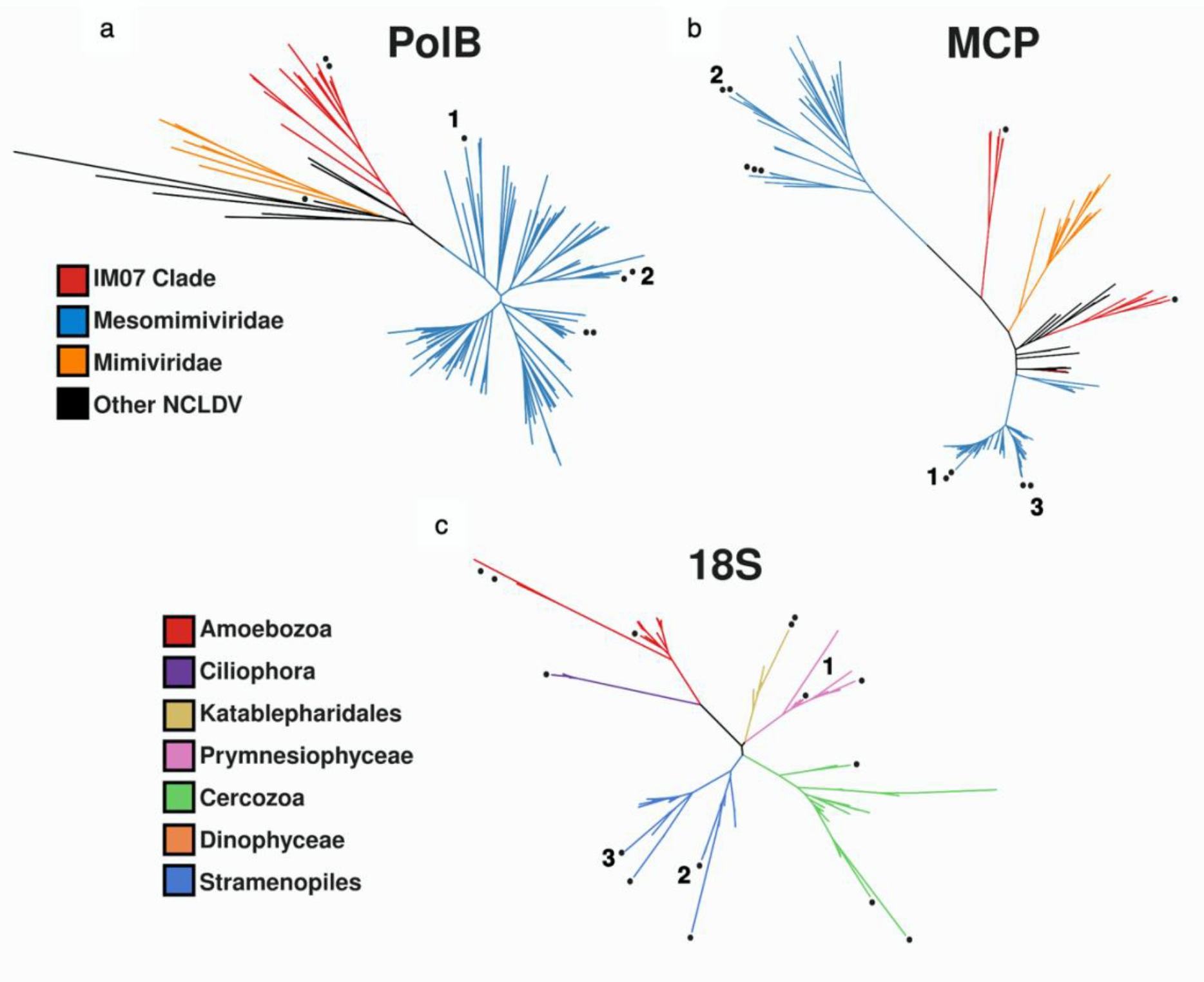
Phylogenetic trees of transcripts assembled from host-virus pairs. Points denote transcripts assembled from single-cell transcriptomes. Numbers denote cells for which transcripts are present in both viral and host trees. (a) DNA-Polymerase subunit B (virus) (b) Major capsid Protein (virus) (c) 18S rRNA (host).

**Extended Data Figure 2.**
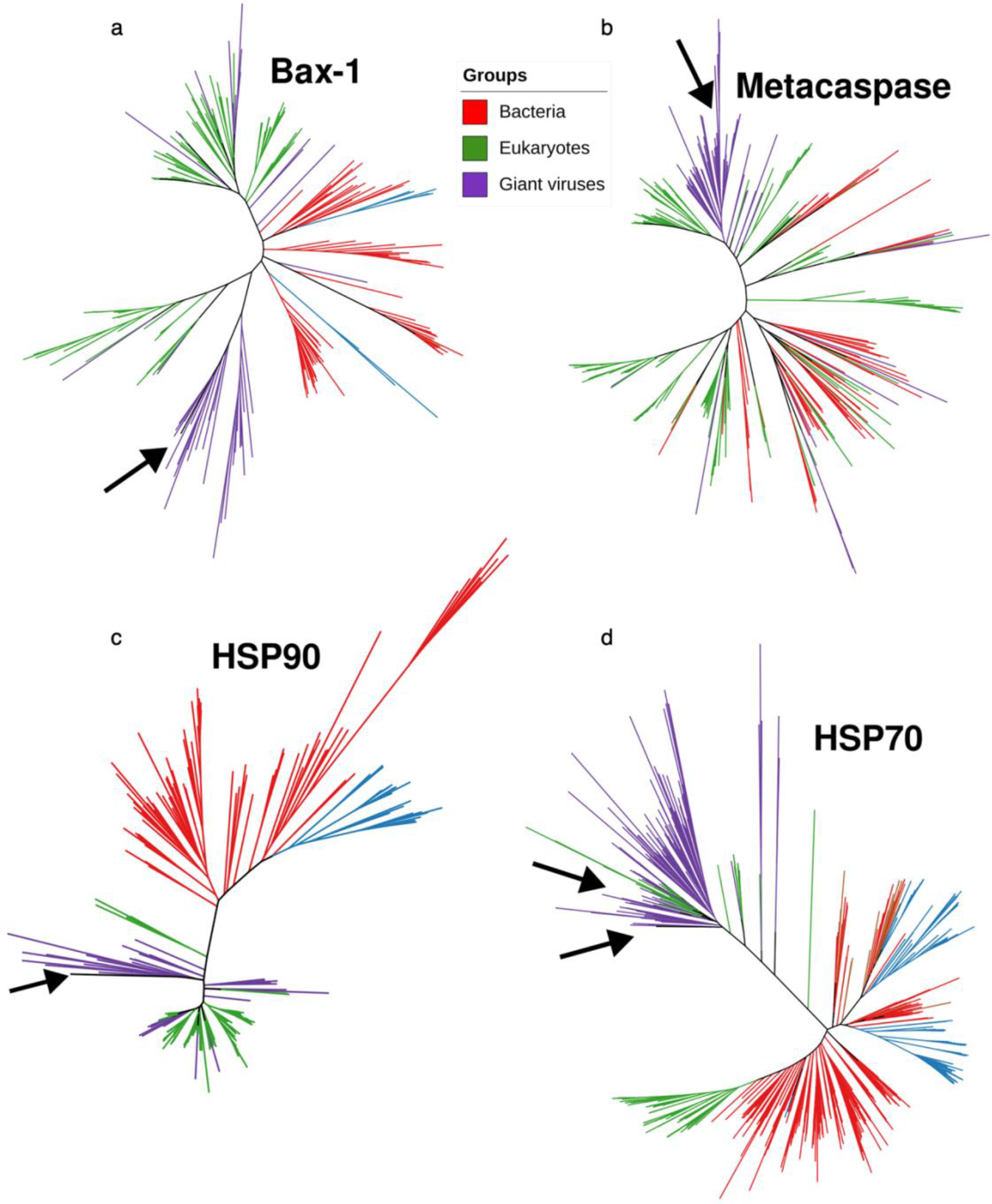
Phylogenetic trees of functional genes present in the predicted *Leucocryptos* virus. The different colors represent bacteria (red), eukaryotes (green), or giant viruses (purple). Arrows point at the location of the predicted *Leucocryptos* virus genes. (a) Bax-1 apoptosis inhibitor (b) Metacaspase (c) heat-shock protein 90 (d) heat-shock protein 70.

**Extended Data Figure 3.**
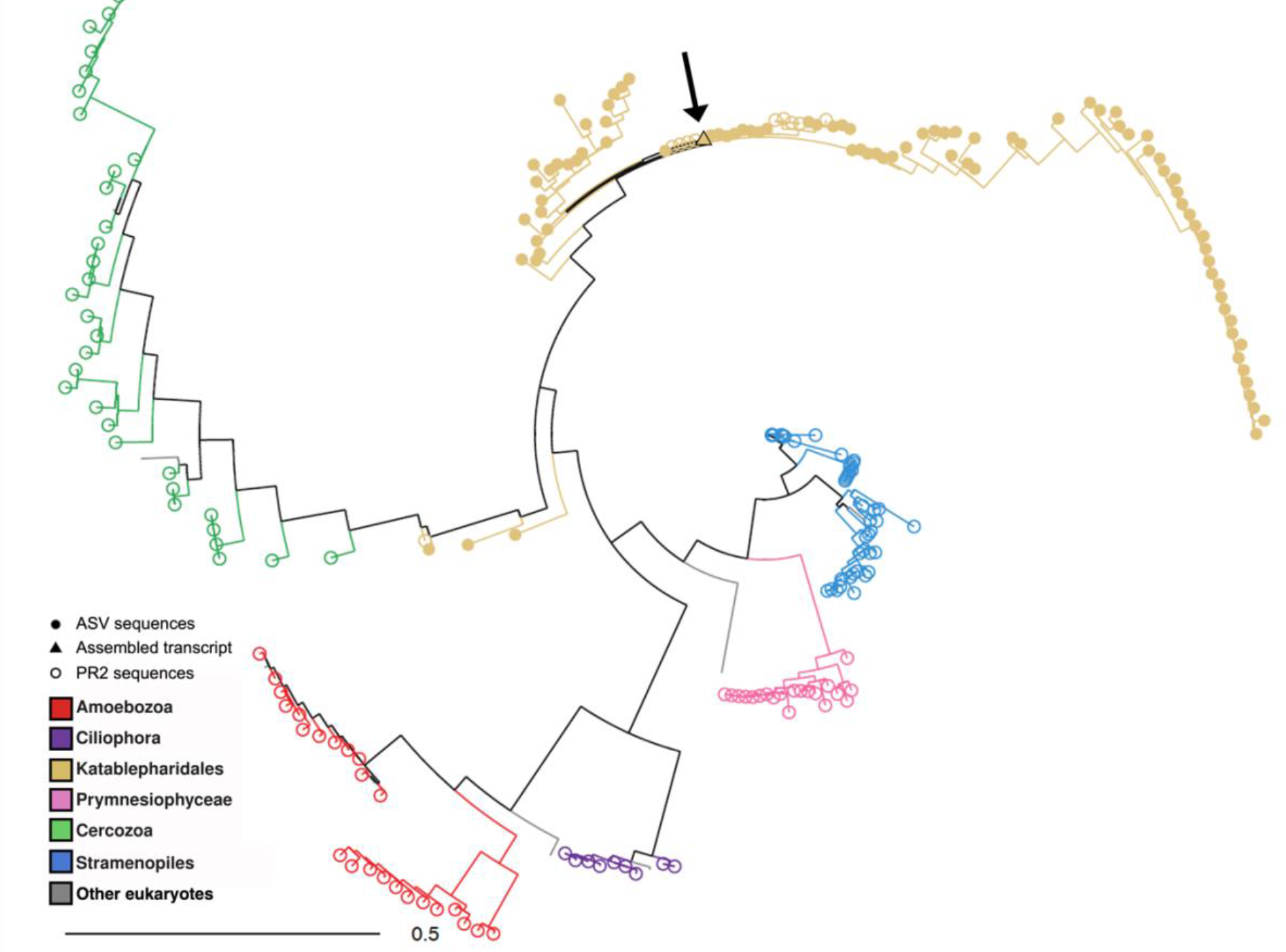
Phylogenetic tree of Katablepharidaceae ASVs, 18S rRNA sequences from PR2 database, and single-cell assembled *Leucocryptos* 18S rRNA gene. The different colors represent the different taxonomic groups analyzed. Filled dots denote ASV sequences while empty dots denote PR2 sequences. The arrow points at the location of the single-cell assembled *Leucocryptos* 18S rRNA gene (in a triangle).

## Supplementary Information

Supplementary Data 1: The homology search results of single-cell reads mapped against the viral gene markers database.

Supplementary Data 2: A summary table of host-virus pairs (as seen in Fig. 2)

Supplementary Data 3: Blast results of infected Katablepharidaceae subpopulation against the PR2 database and against the viral genes markers database.

Supplementary Data 4: Genomic features, gene annotations, and gene expression of the virus GVMAG-M-3300020187-27.

